# Global shortfalls in extinction risk assessments for endemic flora

**DOI:** 10.1101/2020.03.12.984559

**Authors:** R. V. Gallagher, S. Allen, M. C. Rivers, A. P. Allen, N. Butt, D. Keith, T. D. Auld, B. J. Enquist, I. J. Wright, H. P. Possingham, S. Espinosa-Ruiz, N. Dimitrova, J. C. O. Mifsud, V. M. Adams

## Abstract

The Global Strategy for Plant Conservation (GSPC) ambitiously calls for an assessment of extinction risk for all recognised plant taxa by 2020^1^. It is now clear that this target will not be met in the short-term; only 21-26% of known plant species have been assessed^2^ – a monumental shortfall in anticipated knowledge. Yet the need for risk assessments has never been more urgent. Plants are rapidly going extinct^3,4^ and face threats such as climate change^5^ and permanent deforestation^6^. Extinction risk assessments continue to provide the critical foundation to inform protection, management and recovery of plant species^7,8^, the loss of which will have clear consequences for maintaining planetary systems and human well-being^9^. Here, we rank countries of the world based on progress towards assessing the extinction risk to their endemic flora. Overall, 67% of country-based endemic species do not have an extinction risk assessment completed (143,294 species). We show that some of the world’s wealthiest nations, which also have relatively strong species protections, are failing to protect their unique flora by not systematically assessing risks to their endemic species.

---

Target 2 of the GSPC seeks to provide ‘An assessment of the conservation status of all known plant species, as far as possible, to guide conservation action’^1^. To meet this challenge, we argue that initially, the endemic flora of individual countries must be systemically assessed. This declaration requires an objective evaluation of the current performance of countries with regard to completing assessments. To achieve this, we have combined country-level information on five elements: (1) the proportion of endemic plant species with an extinction risk assessment according to the ThreatSearch^10^ database, (2) economic wealth, (3) species protection, (4) population density, and (5) exposure to two key threats: climate warming and permanent deforestation. We use this data to objectively identify countries that may require significant assistance to assess their flora due to low economic status, a lack of species protection or urgent threats.

Plant endemism was estimated across 177 countries and land masses (hereafter ‘countries’) used in the World Geographical Scheme for Recording Plant Distributions (WGSRPD) and accessed via the Plants of the World Online (POWO) database^11^ (Table S1; see Methods and Supplementary Information). While regional differences in taxonomy and risk assessments may exist^12^, standardisation of names between POWO and ThreatSearch ensures an objective comparison across all countries. We identified 215,206 country-level endemic species (Fig. 1) from a total of 331,718 species.

**Figure 1.**
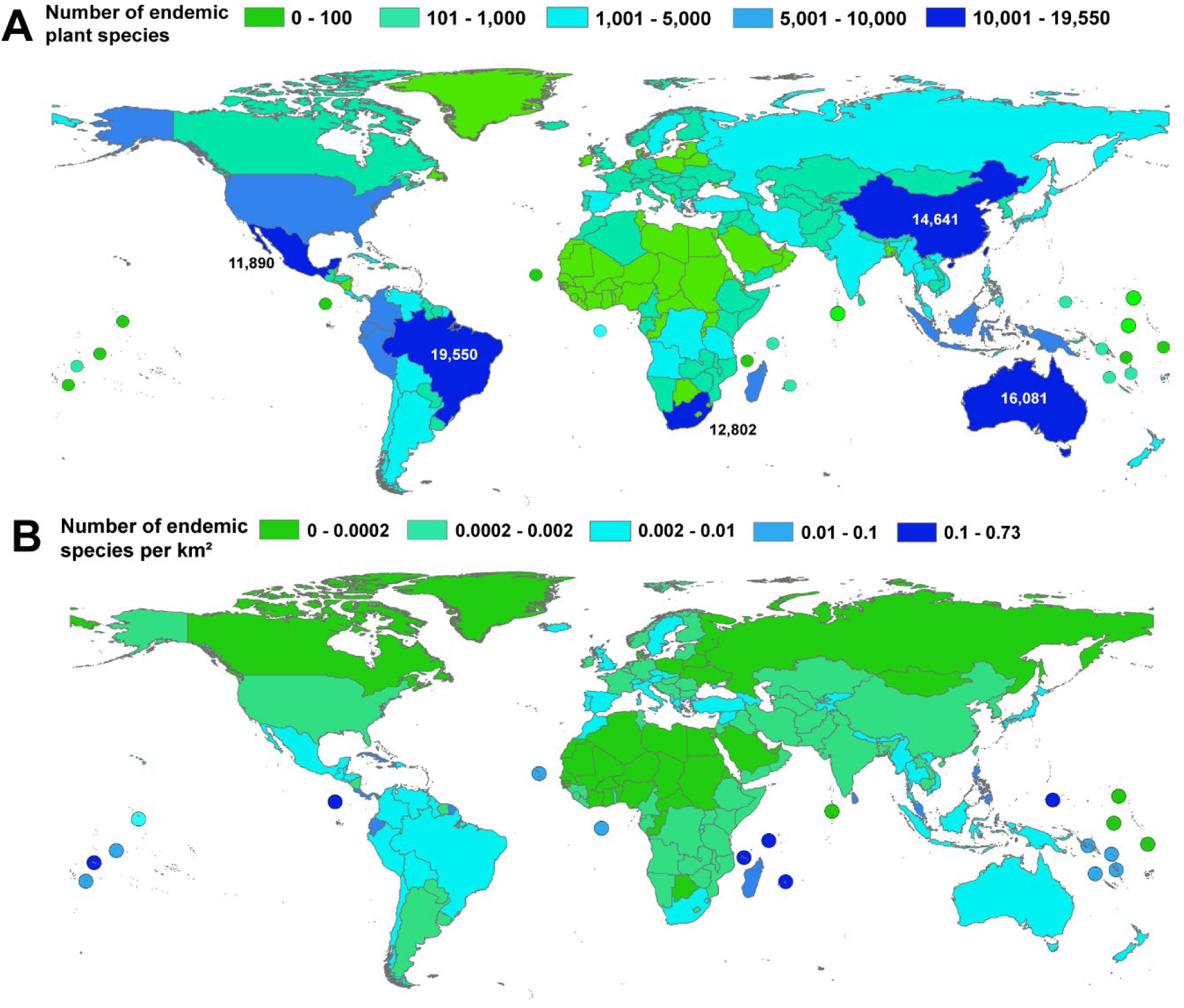
Plant endemism by country based on 1,024,271 records of plant distribution in the Plants of the World Online (POWO) database. Country borders were defined using a modified version of the standardised set of Level 3 spatial polygons developed for the World Geographical Scheme for Recording Plant Distributions (WGSRPD). See Methods and Supplementary Information for detailed explanation and code. (A) Number of endemic plant species in each country. Five countries which have >10,000 endemic plant species are annotated. (B) The number of endemic plant species per km^2^ in each country.

Endemic plant species are unevenly distributed globally, as previously shown^13-15^. Five countries hold the majority of endemic species; Australia, Brazil, China, Mexico, South Africa each provide habitat for >10,000 endemic plant species, which in total represents 35% of all endemic species in our national scale analysis (*n* = 74,964 of 215,206 species). Yet the proportions of endemic species with extinction risk assessed in these five countries varies markedly, from relatively high (0.68 China; 0.85 South Africa) to notably low (0.16 Mexico; 0.22 Brazil; 0.28 Australia) (Fig. 2A). Importantly, high numbers of completed threat assessment in China and South Africa demonstrate that having large numbers of endemic species need not be an impediment to quantifying threat^16^.

**Figure 2.**
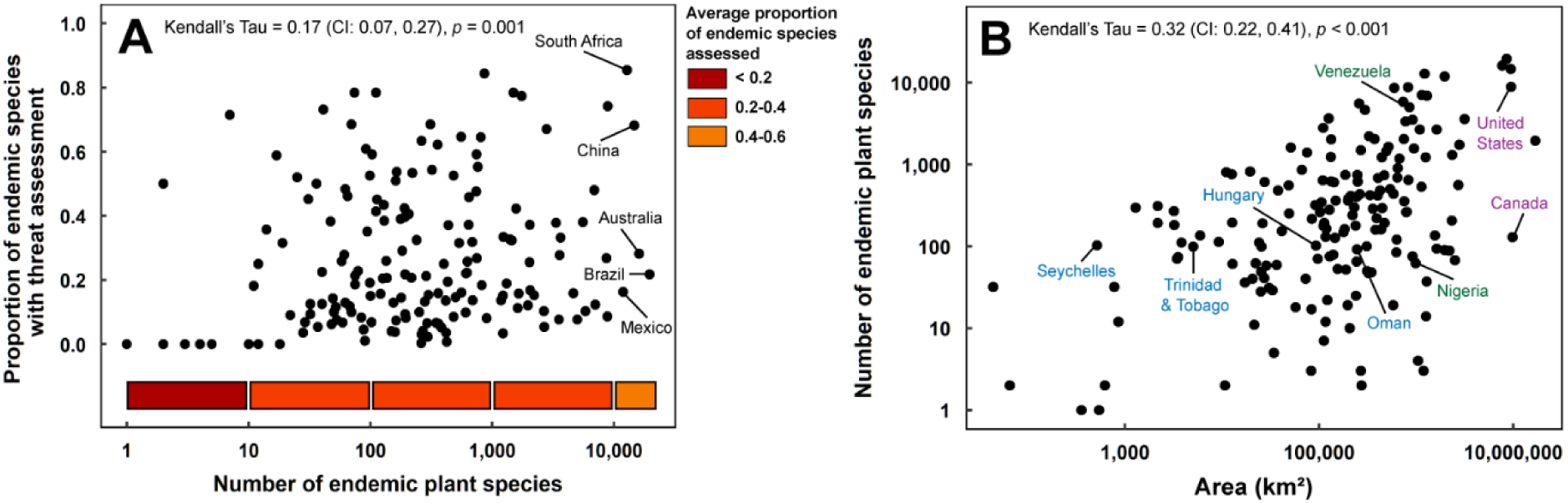
(A) Proportion of endemic plant species per country with an extinction risk assessment in the Threat Search database relative to the size of its endemic flora. Coloured bands below the *x*-axis indicate the average proportion of assessed endemic species in each interval. Countries with >10,000 endemic species are annotated. (B) Number of endemic plant species and area (km^2^) of 177 countries. Annotated countries of the same colour share similar levels of endemism but vastly different area, or vice versa.

As expected, larger countries, on average, have higher numbers of endemic plant species (Fig. 2B). Yet similarly sized countries can support vastly different numbers of endemics (*e.g.*, Canada and the United States (129 vs 8884 endemics respectively and both approximately 9.5 million km^2^); Nigeria and Venezuela (75 vs 3529 endemics; c. 900,000 km^2^). Conversely, countries of different sizes may have similar numbers of endemic plants (*e.g.*, Hungary (92,635 km^2^), Oman (309,669 km^2^), the Seychelles (514 km^2^) and Trinidad and Tobago (5,050 km^2^) all have c. 100 endemics). In many cases high endemism in smaller countries is due to insularity; island countries have an order of magnitude more endemic plant species per km^2^ than do continental countries (ANOVA: F_1,170_ = 17.04, *p* < 0.001; mean: island = 0.04, continental = 0.003) and therefore, in combination with the threats many islands face, they should remain an important focus for plant conservation^13^.

Globally, the proportion of assessed endemic species in a country increases with the total number of endemics, though this relationship is weak (Kendall’s τ = 0.17 (CI: 0.07-0.27); *p* = 0.001; Fig. 2). In countries with relatively modest numbers of endemic species (i.e., < 100; *n* = 64; Fig. 1, 2) assessment rates are, on average, low (22%; range: 0-78%). Some countries in this group are achieving high rates of assessment (e.g. > 70% of species have been assessed in Burundi and Cabo Verde), although fourteen of these countries have assessments for only ≤1% of their endemic plant species. This list includes countries which have endemics from a range of different genera, such as Saudi Arabia (*n* = 90 species in 46 genera; 1 species assessed) and Togo (*n* = 18 species in 15 genera; no species assessed), although it also includes countries with endemic species restricted to a few key genera prone to debates about taxonomic arrangement.

For example, *Rubus, Taraxacum* and *Hieracium* species occur in large numbers in high latitude countries in the Northern Hemisphere, such as Finland, Iceland, Norway and Sweden (Dataset S1). Species in these genera typically reproduce via clones or apomixis and readily hybridise^17-19^, leading to contested taxonomic status, which will need to be resolved prior to threat assessment^20^ and may lower the total number of endemics needing attention in these countries. Regional taxonomic opinion may also differ from the naming conventions used in POWO and the dynamism of systematics can contribute to contested taxonomy between countries.

Although the total number of endemic species is a poor predictor of the number of threat assessments completed by countries (Fig. 2), we hypothesised that economic prosperity and commitment to the protection of species would be positively associated with threat assessment rates globally. That is, risk assessment should be more prevalent in wealthier countries where ecological field studies^21^ and biodiversity data^22^ are often concentrated, or commitment to prioritising conservation may be stronger. However, we found no evidence of a strong relationship between the rate of threat assessment of endemic plants and any of (i) per capita gross domestic product (GDP per capita; Fig 3B), (ii) purchasing power parity with the $USD (Fig. 3C) or (iii) the Species Protection Index (SPI; Fig 3D) which is based on reservation of species ranges in protected areas^23^. Our generalised additive mixed model (GAMM) showed that no deviance could be explained by GDP, PPP, or SPI when accounting for non-linear patterns with a spatial spline smoother (adjusted *R*^2^ = 0.29; deviance explained = 40%; deviance explained by non-linear predictors = 0%; Table S1-3).

**Figure 3.**
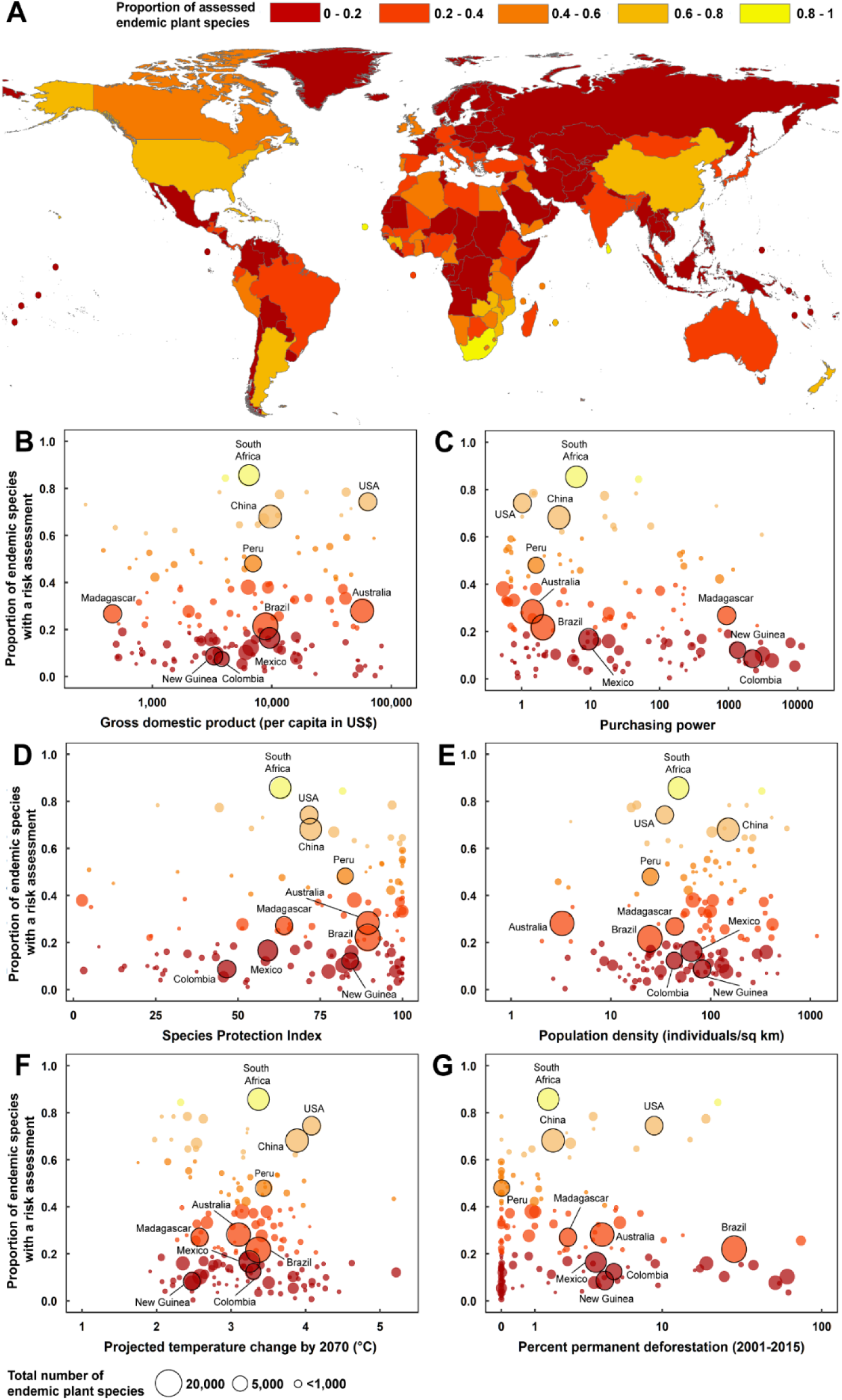
(A) Proportion of endemic species in each country with a threat assessment based on combining the Plants of the World Online (Royal Botanic Gardens, Kew) and Threat Search databases (Botanic Gardens Conservation International; Dataset S2 and Supplementary Information). Associations between the proportion of unassessed endemics in each country and seven predictors are shown: gross domestic product per capita (B), purchasing power parity with the $USD(C), Species Protection Index (D), human population density (E), exposure to change in temperature by 2070 (F), and % permanent deforestation in forests with >30% canopy cover according to^6^ (G). Circle sizes in (B-G) are proportional to the total number of endemic species in each country and the ten countries with the highest number of endemics globally are annotated.

Our statistical model also included a measure of human population density (individuals per km^2^; Fig. 3E) and two metrics of threat exposure for plants (% of permanent deforestation in forests with a >30% canopy cover, and average projected change in mean annual temperature by 2070; Fig. 3F-G). We expected higher population density to be associated with higher proportions of assessed species due to a greater concentration of people in the landscape but found little effect (*p* = 0.08; Table S2). In addition, we tested whether higher threat exposure was reflected in an increased number of completed assessments. This hypothesis was not supported (MAT change: *p* = 0.57; permanent deforestation: *p* = 0.06; Table S2). Therefore, progress toward achieving threat assessment targets for plants is conspicuously uncoupled from economic prosperity, population density, commitment to species protection, and threats.

Our analysis can be used to encourage governments of countries with significant risk assessment shortfalls and large numbers of endemic plants to complete assessments and attract international donors and funding. For instance, countries such as Bolivia, Vietnam and Iran each hold approximately 1% of the world’s endemic plant species (*n* = 7,541 species in total; Fig. 4). These three countries have relatively lower per capita wealth, but Iran also has only marginal levels of species protection, as well as being at high risk from warming temperatures (Fig. 4). Similarly, Madagascar – with approximately 4% of all endemic plant species globally (*n* = 8,640 species) – is clearly identified as a priority country for assessments by our analysis. Only 27% of Madagascan plant species have an extinction risk assessment and this country has economic challenges (Fig. 3-4). Significant deforestation is occurring in Madagascar^24^, though this is often attributed to clearing for shifting agriculture rather than permanent deforestation (i.e., urbanization or commodity driven agriculture)^6^. It is currently unknown how these distinctions between forest loss types may affect extinction risk. South-east Asia is clearly a hotspot for the loss of closed canopy forests (i.e. forests with cover >30%^6^), with several countries in the region at high risk (e.g. Philippines, Borneo, Indonesia, Vietnam).

**Figure 4.**
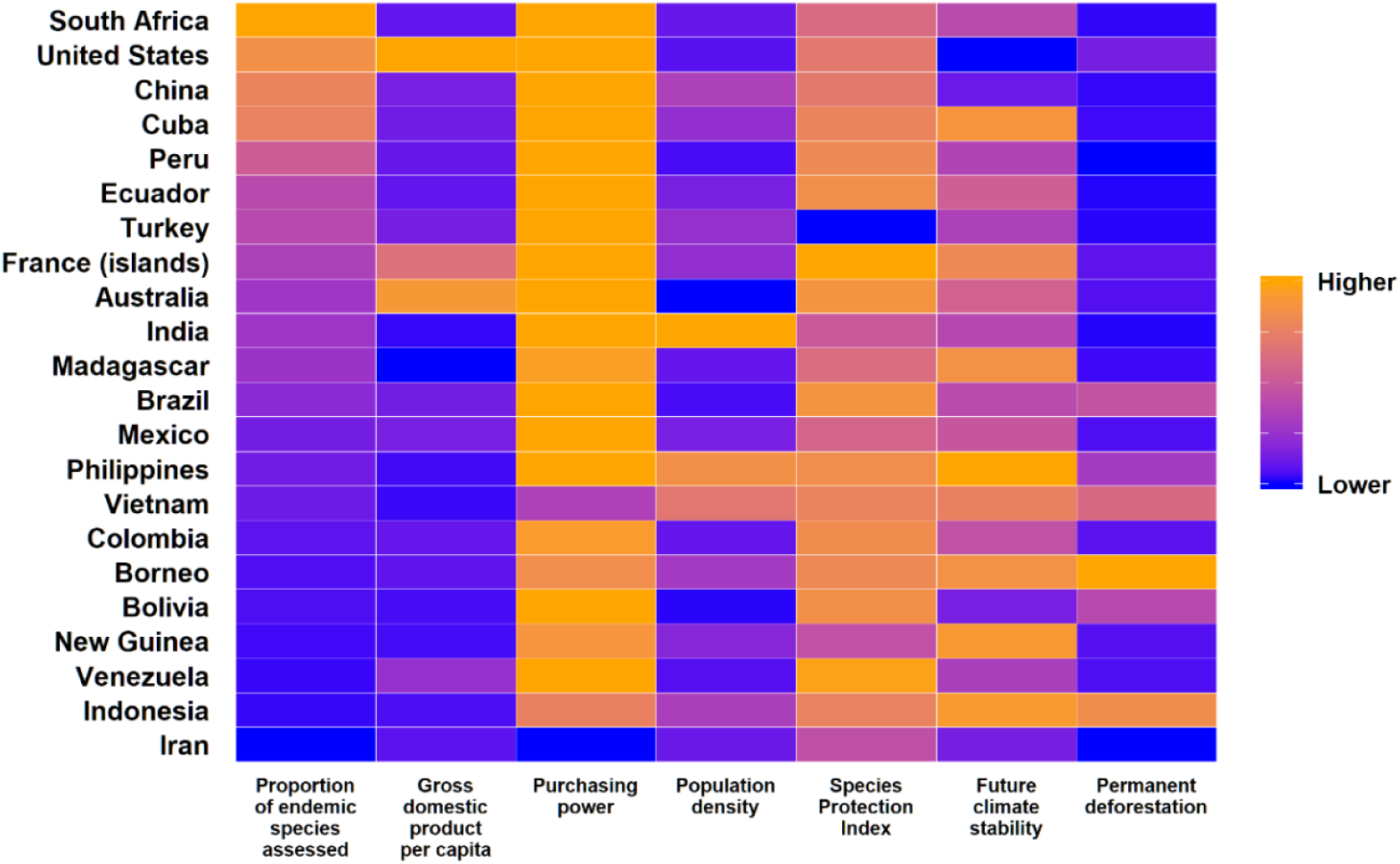
Heat map organising countries based on seven elements: completeness of species assessments, economics (GDP per capita, purchasing power parity), population density, species protection and, threat exposure (climate change and permanent deforestation in forests with >30% canopy cover). The countries displayed each contain >1% of the world’s endemic plant species and are displayed in decreasing order according to the proportion of species assessed for extinction risk.

Clearly, if strategies for management and conservation of the world’s plants are to be well informed, new approaches are needed to address the large global shortfall in risk assessments. Our analysis shows that 143,294 country-level endemic plant species currently lack extinction risk assessment. Individual governments need to play a much larger role in protecting plant species that exist within their borders, starting with an increased commitment to the process of assessment. We do not intend that being endemic should immediately qualify a species as extinction prone – this is an oversimplification. Several large countries have endemic taxa with very large range sizes, such as Australia where 88% of species are endemic according to POWO but range size is, on average, 235,829 km^2^ and varies between 100-7,114,754 km^2,25^. To identify at-risk species, prioritisation based on sound ecological principles and threat exposure will be required in hyper-diverse countries. This may include targeting taxa which are phylogenetically or functionally distinct^26^ or focusing on national areas of high rarity^27^ or endemism^28,29^ such as known ‘biodiversity hotspots’. Rapid or cursory assessments will be less useful than assessments against IUCN Red List criteria which provides extended benefits well beyond determining a species risk status. These include the understanding of population structure, size, and fluctuations, and threats, which are key to well-informed conservation strategies.

The expertise required for conducting risk assessments is likely to be unevenly distributed across the globe. For instance, approximately 90% of ecological field studies are already conducted in objectively wealthy nations^21^ (e.g. United States, Japan, Germany) and geographic biases exist in the spread of taxonomic expertise^30^. This clustering of scientific expertise emphasises the need to support the transfer of skills and emerging technologies^31,32^ to countries with low assessment rates. Respectful and genuine engagement with local and traditional ecological knowledge will be essential for accurately documenting the distribution and ecology of species under assessment.

Several bodies in plant conservation (*e.g*., International Union for Conservation of Nature (IUCN), Botanic Gardens Conservation International, Royal Botanic Gardens Kew) are already training and supporting teams of experienced assessors in countries with high endemism. For instance, specialist workshops targeting known gaps in families such as Myrtaceae and Proteaceae have recently been hosted in Australia. This strategy for completing assessments will undoubtedly translate to a reduction in the shortfall in threat assessments. However, Australia is a relatively wealthy nation and has legislative measures for conducting threat assessments and enforceable species protections. We argue that governments should seek to work closely with international conservation partners to independently assess and subsequently protect their endemic flora, following the lead of nations such as South Africa and China, that have already made plant biodiversity a priority.

Although a considerable number of endemic plant species lack threat assessments, we must redouble efforts to meet the ambitious goals of the GSPC. With the addition of non-endemic taxa to these efforts funding for conserving plant diversity globally needs to immediately increase. Shortfalls in assessments translate into knowledge gaps for planning and management of endemic species – a major and urgent issue. Assessing species to identify levels of threat is a critical first step to allow subsequent listing of species on statutory threatened species lists, trigger species recovery planning, and subsequent design of management and monitoring strategies for threatened species to safeguard against extinction and facilitate recovery. In addition, when completing threat assessments, we expand our knowledge of the natural world through the systematic documentation of species’ taxonomy, biology, ecology and demography. The assessment process compels cross-disciplinary experts to work collectively toward a stronger understanding of the diversity of life on Earth, to all our benefits.

## METHODS

### Countries

The world was divided into 177 modified countries (hereafter ‘countries’) based on the Level-3 spatial units in the World Geographical Scheme for Recording Plant Distributions^33^ (WGSRPD; Fig. S1-2; Dataset S3 and S4). These countries largely correspond to accepted political boundaries (*i.e., n* = 97 of 177 countries match political boundaries; Fig. S3). Several spatial aggregation procedures were used to create the remaining 79 countries. These procedures harmonised the boundaries of WGSRPD units to the borders of sovereign nations (Figs. S4-8; see Supplementary Information for full methods). For instance, Australia is an aggregate of the seven mainland states and six offshore territories that it governs (*i.e.*, Christmas, Cocos (Keeling), Heard, Macquarie, McDonald, and Norfolk Islands); Fig. S4). Other countries represent WGSRPD units which contain multiple sovereign nations (*e.g.*, the Baltic States of Estonia, Lithuania, Latvia were analysed as one country, as was the island of Borneo which is claimed by Malaysia, Indonesia and Brunei; Fig. S6). As data on endemic species richness was not available at finer resolution, these multi-nation countries could not be disaggregated further. Note that several countries (*e.g.*, French offshore islands) could not be aggregated with their sovereign nation (here, France) because the sovereign itself was part of a multiple-nation country (i.e., the level-3 unit containing continental France also contains the sovereign nation of Monaco and the British territory of Jersey; Fig. S7). Antarctica and the disputed regions of Gaza, Jamu-Kashmir and Kosovo were excluded from analyses (Fig. S8).

### Endemic species data and threat assessments

Counts of the number of endemic species in each country were based on data from the Plants of the Word Online (POWO) database that is maintained by Royal Botanic Gardens, Kew^11^. We assembled an initial dataset of 1,024,271 records of plant occurrence across 373 WGSRPD Level-3 units (*n* = 331,718 species). Species were considered endemic to a country if they occurred solely within its borders (*n* = 215,206 endemic species globally; Fig. 1; Dataset S2). We excluded from our analyses all taxa with unaccepted names (e.g. synonyms or unresolved names), as well as those considered introduced, extinct or to have a doubtful location according to POWO. The POWO dataset represents a wide breadth of known plant diversity, including species from 452 families (i.e., angiosperms (*n* = 404 families), gymnosperms (*n* = 12 families), pteridophytes (*n* = 45 families), bryophytes (*n* = 1 families); Fig. S9).

The ThreatSearch database^10^ maintained by Botanic Gardens Conservation International was used to determine what proportion of the 215,206 endemic plant species have an assessment of their extinction risk (Fig. 2; S10). Assessments listed in ThreatSearch vary in scope from global to regional (e.g. within individual states of a country, or across multiple countries and not all are based on IUCN Red Listing Criteria^34^; Dataset S5). Previous matching between POWO and ThreatSearch mean they are readily comparable, though their naming conventions may differ from regional taxonomic opinion. We considered a species to have been assessed if ThreatSearch documented any record for the taxon, including its subspecies or varieties (e.g., 5.8% of species were assessed as infraspecific ranks). Analyses were performed in R using the *tidyverse*^35^ collection of packages (see Dataset S6 for analysis code).

### Economic metrics and population density

Gross Domestic Product per capita (GDP) in $USD, Purchasing Power Parity conversion factor (PPP) and Total Population data were gathered from the World Bank^36^ and World Fact Book^37^ and appended to each country. GDP per capita reflects the perception of wealth in a nation’s people; this may relate more strongly to willingness for greater environmental protection than raw GDP. PPP estimates the units of a country’s currency needed to buy the same amounts of goods and services in the domestic market as a US dollar would buy in the United States. Total population was converted to population density using estimates of land area per country derived from a Behrmann Equal Area projection. In countries which incorporate more than one sovereign nation (e.g. Borneo, the Baltic States; Figs. S4-8), variables were weighted by the land area of contributing countries. For example, GDP per capita/PPP/population density for the island of Borneo is derived from values for Indonesia, Malaysia and Brunei Darussalam (see Supplementary Information for full methods for calculation, Dataset S6 for analysis code and Dataset S2 for data).

### Species protection index

The 2018 Environmental Performance Index ranks countries based on 24 metrics of environmental and ecosystem health^23^, including a Species Protection Index (SPI). The SPI estimates the average proportion of species’ distributions in a country within protected areas. SPI values were appended to each country, using weightings where needed (see Economic metrics and Supplementary Information for full methods for calculation and Dataset S6 for analysis code).

### Climate warming exposure

Exposure of each country to climate warming was estimated using temperature anomalies between current climate and future climate projections. Anomalies were produced by subtracting gridded global data on current (1979-2013) mean annual temperature conditions (MAT; °C) from a median projection of MAT conditions derived from seven global climate models (GCMs) for the decade centred on 2070 (2061-2080) under representative concentration pathway 8.5. (Fig. S11A) Current climate data and projections for GCMs (*i.e.*, CESM1-BGC, MPI-ESM-MR, MIROC5, CMCC-CM, CESM1-CAM5, IPSL-CM5A-MR, FIO-ESM) were sourced from CHELSA at a 30 arc-second resolution^38^. GCMs were selected to minimise interdependencies in model creation^39^. The average (mean) anomaly in MAT conditions across each country was then used to quantify potential exposure to climate change (Fig. S11B). In this analysis, temperature warming is used as a generic proxy for other climate-related threats. We assume that averaged anomalies represent basic trends, but do not capture extreme events and that spatial averaging does not systematically bias the projected magnitude of climate warming across countries of different sizes.

### Permanent deforestation

Global maps of the drivers of forest loss^6^ were used to calculate the percentage of forested areas with >30% cover subject to permanent deforestation between 2001-2015 in each country. Permanent deforestation was defined as land use conversion that prevents subsequent forest regrowth^6^. Two drivers of forest loss were combined spatially to identify areas of permanent deforestation: commodity-driven deforestation (category 1 in^6^) and urbanization (category 5 in^6^) (Fig. S12). This composite map is based on classification of remotely sensed imagery used to identify all 10 km x 10 km grid cells (Berhmann equal-area projection) where drivers of permanent deforestation were the most likely cause of forest disturbance since 2000. Areas with canopy cover <30% will not be captured by this mapping and consequently deforestation is likely to be underestimated in semi-arid and some cold climates.

### Statistical analyses

The Kendall rank correlation coefficient (Kendall’s τ) was used to test for an association between (i) the number of endemic species and country area (km^2^); and (ii) the proportion of endemic species with a threat assessment and the total number of endemic species in a country (Fig. 2). Confidence intervals for τ were calculated via bootstrapping using the function *KendallTauA* in package *DescTools*^40^ in R^41^. ANOVA was used to test for differences in the number of endemic species per km^2^ between island and continental countries.

Generalised additive mixed models (GAMMs) were used to quantify the relationship between the proportion of endemic species assessed in countries; a set of economic, demographic and threat predictor variables (*i.e.*, GDP per capita, PPP, population density, SPI, climate change exposure and permanent deforestation; Table S1); and the spatial distance between countries (latitude/longitude of country centroids). GAMMs were chosen as they allow the effect of non-linear spatial patterns to be combined with linear predictors. The latitude and longitude coordinates at the centroid of each country were added to GAMMs as a two-dimensional spherical spline smoother, *f*(lat,lon).

All variables, except climate change exposure, were transformed prior to analysis as follows: (1) skewed variables (i.e., GDP per capita, PPP, population density) were log-transformed; (2) bounded variables with few values equal to 0 or 100 (i.e., SPI, proportion of endemics assessed) were logit transformed; (3) bounded variables with many values equal to 0 or 100 (i.e., permanent deforestation; *n* = 64 of 165 observations were true zero’s) were converted to a binary factor (TRUE/FALSE). We assumed a Gaussian distribution and identity link and partitioned variance into spatial and aspatial components (and their covariance) to estimate the deviance explained by linear predictors. Tests of model residuals (Moran’s I) indicated that the addition of the spatial spline adequately accounted for non-linear trends (Table S3). Several routine diagnostic plots confirmed that GAMM assumptions were reasonably met, including checks for normality, equal variance of residuals and linearity (Fig. S13-17). All analyses were implemented via packages *GAMM4*^42^, *ape*^43^, *geosphere*^44^, *gratia*^45^ and *ecospat*^46^.

## ACKNOWLEDGEMENTS

Steven Bachman from Royal Botanic Gardens Kew provided access to data from the Plants of the World Online and valuable feedback on earlier versions of the draft. R.V.G. was supported by Australian Research Council DECRA Fellowship (DE170100208).

## AUTHOR CONTRIBUTIONS

R.V.G conceived the original idea with V.M.A. and wrote the first draft. M.R., A.P.A., N.B., D.K, T.D.A., B.J.E., I.J.W, and H.P.P. extended the ideas and scope of the manuscript and contributed substantively to writing and revisions. R.V.G., M.R., S.A., A.P.A, N.D., S.E.-R., J.C.O.M compiled and analysed the data.

## DATA AVAILABILITY

All data and code for reproducing analyses is available in the Dataset S6 and processes are described in the Supplementary Information.

## COMPETING INTERESTS

The authors declare no competing interests.

## ADDITIONAL INFORMATION

### Supplementary Information

**Dataset S1:** Endemic species per country according to POWO and their extinction risk assessment status according to ThreatSearch

**Dataset S2:** Country-level endemism, assessment completion, economic and environmental metrics, population density, and threats

**Dataset S3:** World Geographical Scheme for Recording Plant Distributions units and corresponding sovereign nations

**Dataset S4:** Spatial layer of countries developed from World Geographical Scheme for Recording Plant Distributions units

**Dataset S5:** Sources within the ThreatSearch database

**Dataset S6:** Analysis code and data

