## Supplementary Material for "Global shortfalls in extinction risk assessments for endemic flora"

### 1. Overview

Target 2 of the Global Strategy for Plant Conservation (GSPC) calls for extinction risk to be assessed for all recognised plant taxa by 2020[1]. Yet only 21-26% of known plant species had been assessed by 2018[2] and the need for risk assessments has never been more urgent[3-7]. One way to address this shortfall is for countries to concentrate on assessing their endemic plant taxa. Here, we rank countries of the world based on progress towards assessing extinction risk in their endemic flora. We combine country-level metrics of economic wealth, species protection status, population density and threat to highlight how these factors are de-coupled from progress towards plant conservation globally. Below, we outline a series of methodological steps that can be used to recreate our analyses. Code and relevant data are available in Dataset S6.

### 2. Deriving a country dataset from plant distribution data

Plant distribution data for 331,718 species was accessed in February 2019 from the Plants of the World Online (POWO) database maintained by Royal Botanic Gardens, Kew. Only species with accepted names in POWO (i.e. no synonyms, or unresolved names) were included in this analysis. POWO documents plant distributions using spatial units (polygons) from the World Geographical Scheme for Recording Plant Distributions (WGSRPD); plant species are recorded as Native, Introduced or Extinct in each spatial unit. We used data from Level-3 WGSRPD spatial units (Fig. S1) as the basis of our analysis.

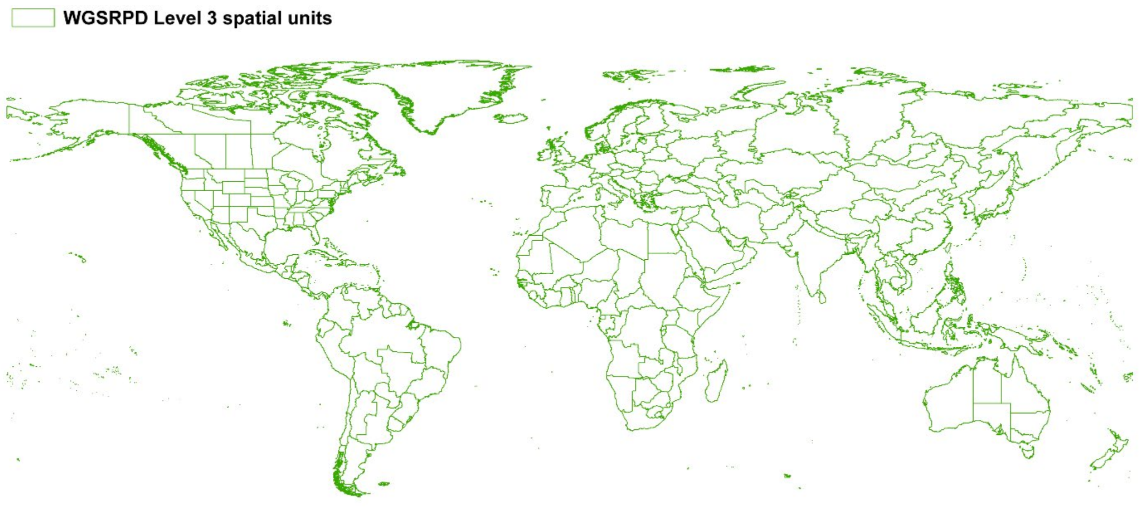


**Figure S1.** Level-3 spatial units World Geographical Scheme for Recording Plant Distributions (WGSRPD) used to record plant distribution information in the Plants of the World Online (POWO) database.

Level-3 WGSRPD units do not always correspond directly to the political boundaries of countries (Fig. S2), though we needed to assign country-level metrics on economics, threats and species protection. To address differences in borders, we created a set of modified countries (hereafter 'countries') that harmonise the boundaries of plant spatial distributions to the borders of sovereign nations. A spatial layer of sovereign nations was accessed from Natural Earth Data[8].

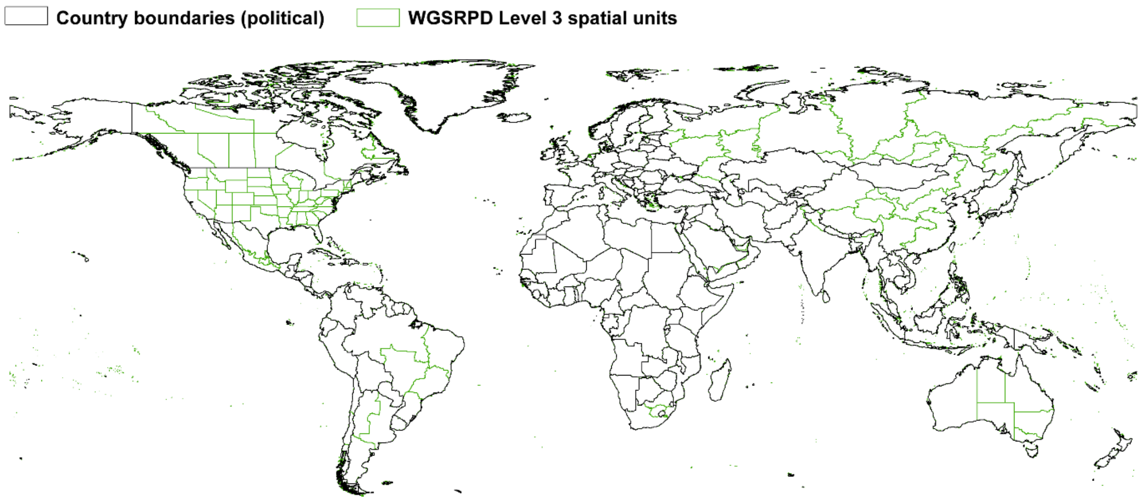


**Figure S2.** Relationship between Level-3 spatial units in the World Geographical Scheme for Recording Plant Distributions (WGSRPD) used to record plant distribution information in the Plants of the World Online (POWO) database (green) and the political boundaries of countries (black).

Several scenarios arose when creating our country polygons, as follows:

**Boundaries of WGSRPD units and sovereign nations were identical (*n* = 97 countries; Fig. S3).** 55% of countries required no spatial analyses to reconcile boundaries between plant distribution data and political boundaries.

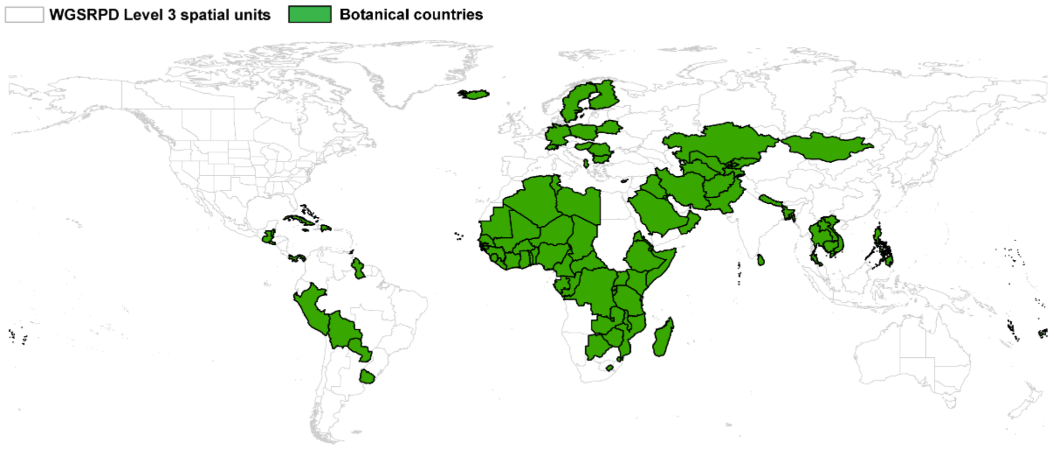


**Figure S3.** Countries whose borders correspond directly with Level-3 spatial units in the World Geographical Scheme for Recording Plant Distributions (green polygons).

**Multiple WGSRPD units were aggregated to represent a single, sovereign nation (*n* = 21 countries).** For instance, the country of Australia is an aggregate of WGSRPD units for seven mainland states and six offshore territories it governs (i.e., Christmas, Cocos (Keeling), Heard, Macquarie, McDonald, and Norfolk Islands; Fig. S4). A similar situation arose for Brazil, which is an aggregate of five WGSRPD units for the regions of Brazil Northeast, Brazil North, Brazil West-Central, Brazil Southeast, and Brazil South.

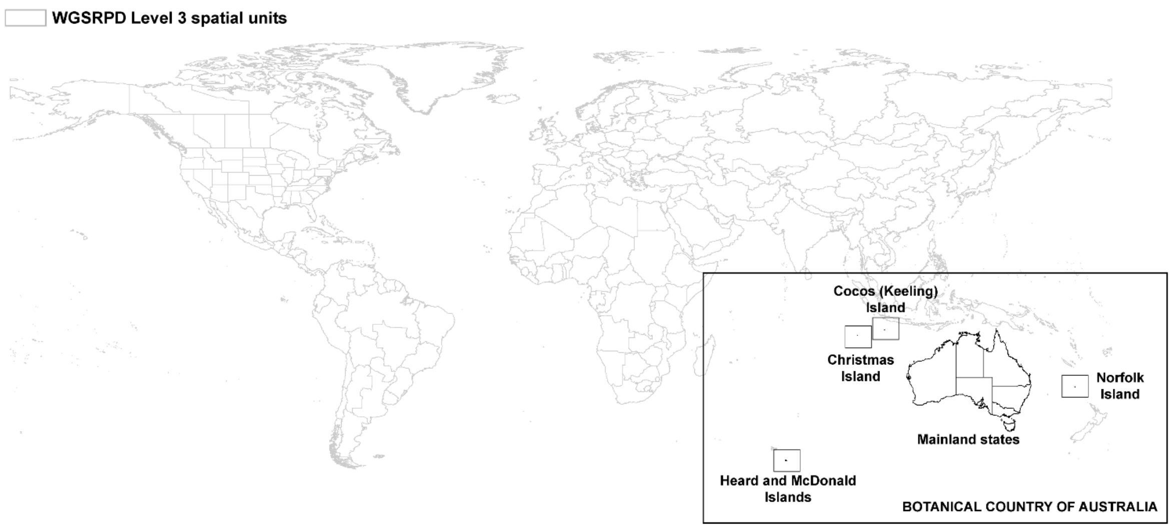


**Figure S4.** The country of Australia in our analysis. Inset map shows the World Geographical Scheme for Recording Plant Distributions (WGSRPD) Level-3 spatial units aggregated to create the country of Australia. All WGSRPD Level-3 spatial units aggregated are part of Australia's sovereign territory.

**A single WGSRPD unit contains all or part of multiple sovereign nations (*n* = 11 countries).** For instance, a single WGSRPD unit contains both Austria and Liechtenstein (Fig. S5). Therefore, endemic species in this spatial unit must be attributed to a country containing two sovereign nations. Countries of this kind must have estimates of Gross Domestic Product per capita (GDP per capita), Purchasing Power Parity (PPP), Species Protection Index (SPI) and population density (individuals/km$^2$) calculated objectively based on the total land area belonging to each sovereign nation (see **5. Economic, protection and population metrics for countries** below).

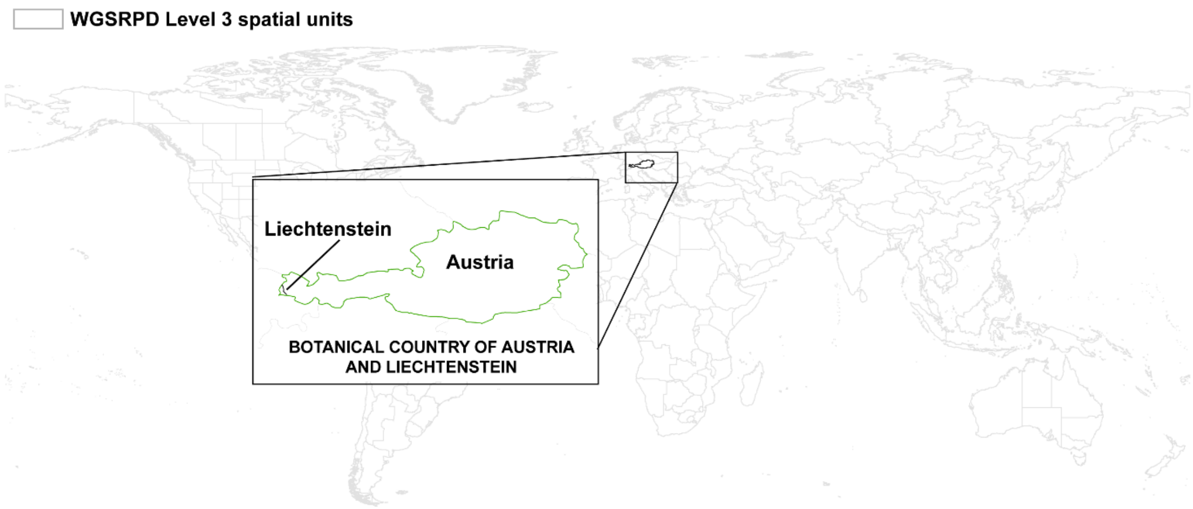


**Figure S5.** The country of Austria and Liechtenstein in our analysis. Inset map shows the World Geographical Scheme for Recording Plant Distributions (WGSRPD) Level-3 spatial unit encompassing both sovereign nations.

A similar situation arises for the island of Borneo which is represented by a single WGSRPD unit but is composed of three sovereign nations: Indonesia (72.5% of land area), Malaysia (26.7% of land area) and Brunei (0.8% of land area) (Fig. S6). However, in this case, the WGSRPD unit for Borneo contains only 28.5% of the sovereign nation of Indonesia and 59.6% of Malaysia, but 100% of Brunei.

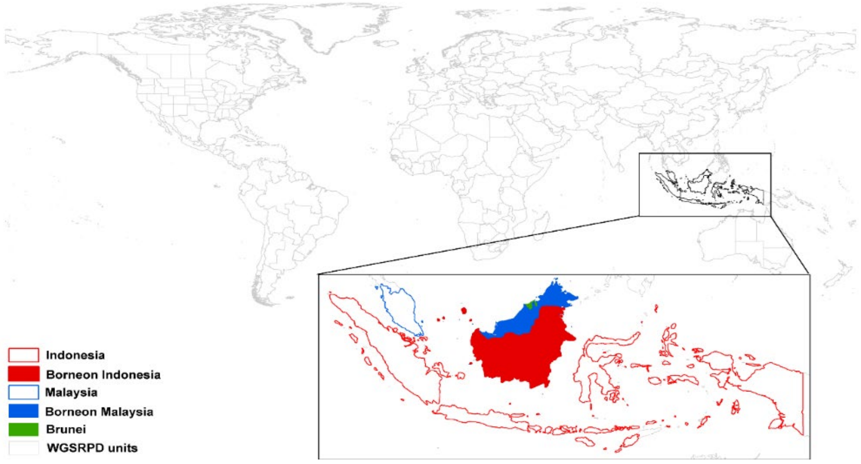


**Figure S6.** The 'country' of Borneo. Inset map shows the World Geographical Scheme for Recording Plant Distributions (WGSRPD) Level-3 spatial unit for Borneo (filled polygon). Borneo is composed of land claimed by three sovereign nations: Indonesia (72.7% of Bornean land area), Malaysia (26.6%) and Brunei (0.08%). Open polygons show land area of Indonesia and Malaysia outside Borneo (71.5% and 40.4% respectively).

**All WGSRPD units belonging to a single sovereign nation cannot be aggregated to a single country (*n* = 23 countries).** For example, France and its offshore territories are two separate countries in our analysis as the WGSRPD unit for continental France also contains the sovereign nation of Monaco and the British island territory of Jersey (Fig. S7). The GDP per capita, PPP, SPI and population for the WGSRPD unit for continental France needs to also incorporate data for Monaco and Great Britain, though all French offshore territories can simply be assigned these metrics for France.

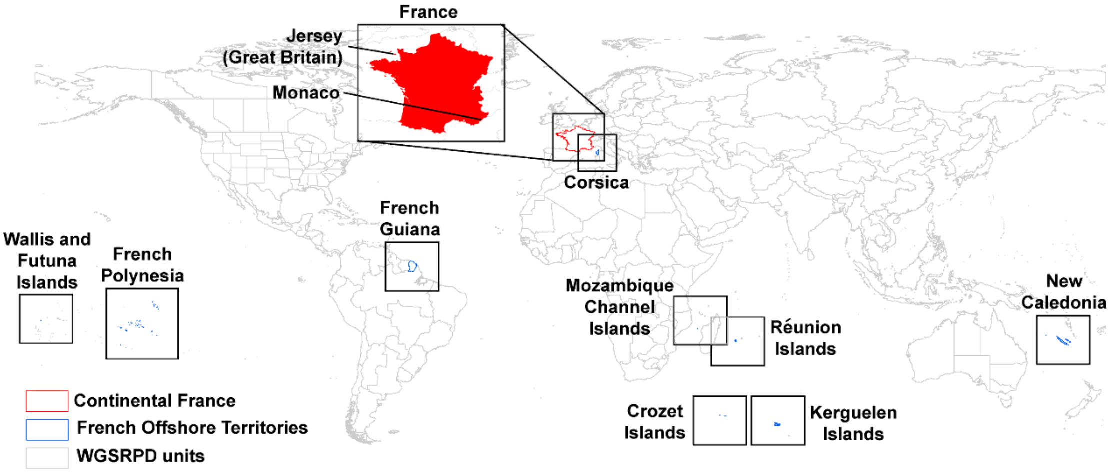


**Figure S7.** The countries of Continental France and French offshore territories. Inset map shows the World Geographical Scheme for Recording Plant Distributions (WGSRPD) Level-3 spatial unit for continental France (filled red polygon), which also includes the sovereign nation of Monaco and the Great Britain territory of Jersey. French offshore territories (open blue polygons) are combined into a single, and separate, country than continental France.

Using these methods, we identified 177 countries globally (Fig. S8). See Dataset S3 for a full list of sovereign nations and how they relate to WGSRPD units. This spatial layer is available as Dataset S4.

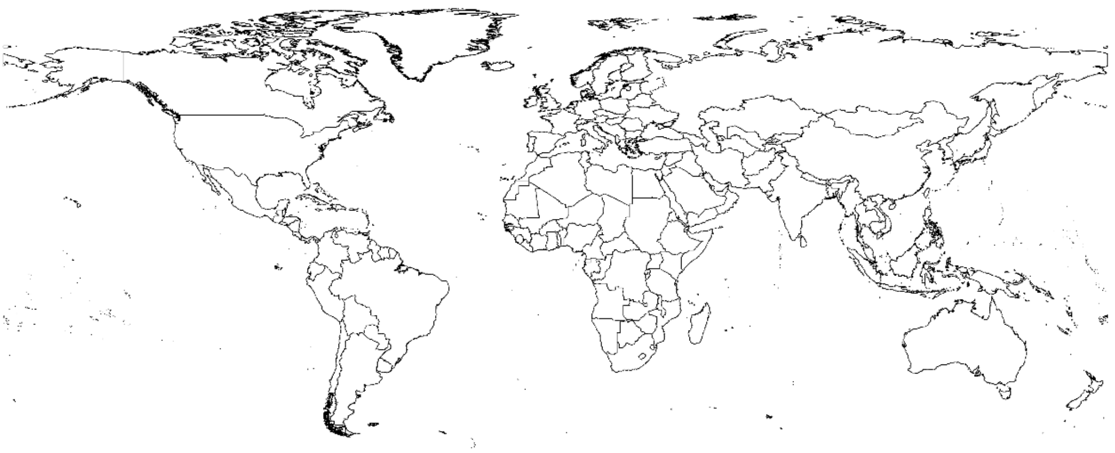

**Figure S8.** Countries of the world based on the aggregation of Level-3 spatial units in the World Geographical Scheme for Recording Plant Distributions (WGSRPD).

### 3. Plant endemism in countries

Using distribution data for 331,718 plant species accessed from the POWO database, we counted and mapped the number of species unique to each country (i.e., endemics; $n$ = 215,206 species). In total, 65% of plant species in POWO as of February 2019 were considered endemic to a single country. Of the 177 countries, 172 have endemic species (i.e., only Kuwait, Marshall Islands, Maldives, Nauru, and Tuvalu have no endemics). The mean and median number of endemics per country were 1216 and 193, respectively.

The POWO dataset represents a substantial portion of known plant diversity, including species from 452 families (i.e., angiosperms (404 families), gymnosperms (12 families), pteridophytes ($n$ = 45 families), bryophytes (1 family); Fig. S9). POWO includes species from regional Floras and monographs, as well as established databases such as Grassbase and PalmWeb. These data sources vary in the extent to which comprehensive synonymy is included, their publication status and their level of peer review; other, more authoritative lists likely exist for specific regions or taxa. POWO is a dynamic resource and we anticipate additions and taxonomic changes will occur. Our analysis code is in Dataset S6 to enable different versions of POWO to be used in re-analysis or new projects. Note that Royal Botanic Gardens, Kew cannot warrant the quality or accuracy of the data. See Dataset S1 for the identity of endemic species per country.

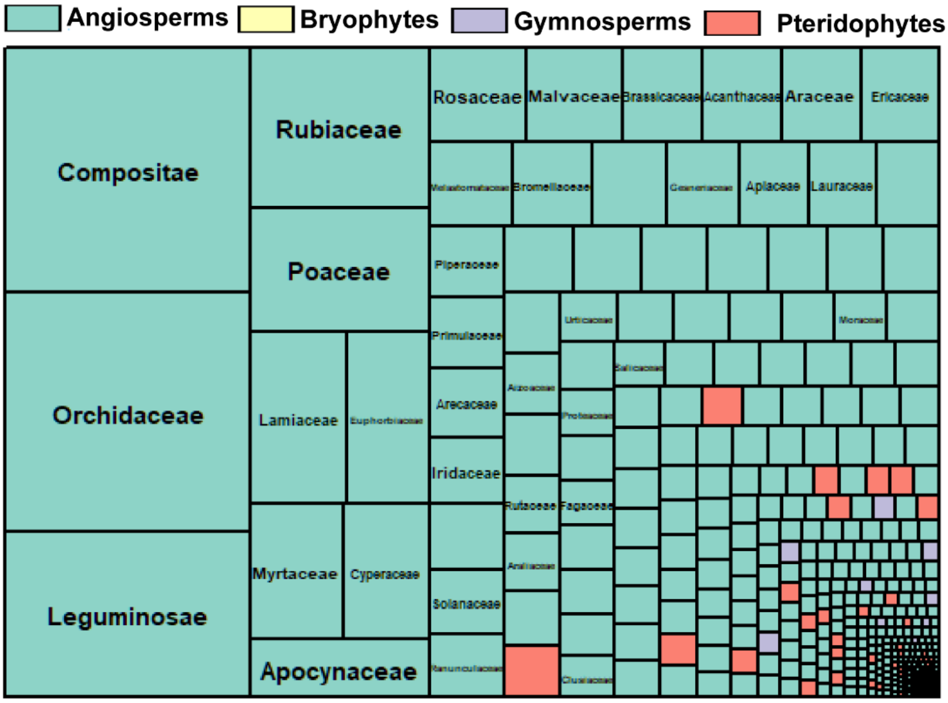


**Figure S9.** Treemap of number of accepted species in Plants of the World Online per family and plant group (Angiosperms, Bryophytes, Gymnosperms, Pteridophytes). Box size is proportional to the number of species per family.

### 4. Extinction risk assessments for countries

Data on the completion of extinction risk assessments for endemic species were accessed from the ThreatSearch database in July 2019. ThreatSearch is maintained by Botanic Gardens Conservation International https://tools.bgci.org/threat_search.php and is the largest global compilation of extinction risk assessments for plants, containing approximately 120,000 assessments. We calculated the proportion of endemic species in each country with an extinction risk assessment by merging data from POWO and ThreatSearch, based on species name (Fig. S10). Previous matching between POWO and ThreatSearch mean they are readily comparable, though these naming conventions may differ from regional taxonomic opinion. We considered a species to have been assessed if ThreatSearch documented any record for the taxon, including its subspecies or varieties (*e.g.*, 5.8% of species were assessed as infraspecific ranks). Of the 215,206 endemic species queried in ThreatSearch, 33% (71,911 species) had an extinction risk assessment (see Dataset S1).

Note that assessments in ThreatSearch vary in scope (*i.e.*, global, regional, national) and many – but not all – use IUCN Red List Criteria to assess extinction risk (see Dataset S5 for a full list of sources). ThreatSearch is a living-resource; new and legacy extinction risk assessments are being added to the database. Therefore, recent versions of the database should be accessed before generating new analyses. All data used in this analysis are available in accompanying datasets; see Dataset S1 for the number of assessed species per country.

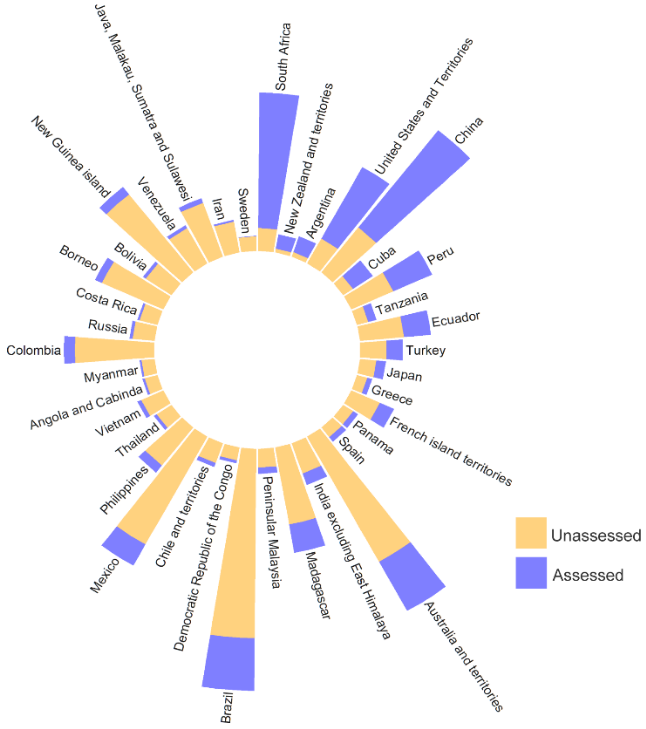


**Figure S10.** Circular bar plot of the proportion of endemic plant species with a extinction risk assessment (blue portion) in the 38 countries of the world with > 1,000 endemic plants.

### 5. Economic, protection and population metrics for countries

For each sovereign nation, data on gross domestic product per capita (GDP per capita), purchasing power parity (a.k.a. PPP conversion factor) and total population were accessed from the World Bank[9] and the CIA World Fact Book[10] and data on Species Protection Index (SPI) was accessed from The Yale Centre for Environmental Law & Policy[11]. PPP conversion factor for GDP per capita is the number of units of a country's currency required to buy the same amounts of goods and services in the domestic market as USD$ would buy in the United States[12]. For all metrics, the most recent year of available data was used. All metrics data can be accessed in Dataset S2.

As described in **2. Deriving a country dataset from plant distribution data**, a country may be shared among several sovereign nations (*e.g.*, the island of Borneo is shared between Malaysia, Indonesia and Brunei) and these metrics must be apportioned relative to land area claimed. Therefore, we intersected a map of WGSRPD units ($n$ = 364) and political boundaries of sovereign nations ($n$ =197) to create a dataset of unique combinations (rows) of WGSRPD and sovereign nations ($n$ = 439). We assigned GDP per capita, PPP, and SPI of the sovereign nation to each row and weighted the sum of these based on the land area of each unit as a proportion of the total area of the country based on equation 1.

Eq.1: $$x^{BOT} = \sum_{i=1}^{n} x_i^{SOV} \left(\frac{a_i^{UNIT}}{a^{BOT}}\right) \mathrm{w}$$

Where $x^{BOT}$ was the estimated GDP per capita/PPP/SPI of the country, $x^{SOV}$ was the GDP per capita/PPP/SPI of the sovereign nation, $a^{UNIT}$ was the area of that sovereign nation within the country, and $a^{BOT}$ was the total area of the country.

For example, the island of Borneo comprises the sovereign nations of Indonesia, Malaysia and Brunei. The GDP per capita, PPP and SPI for these countries were weighted by the total area of Borneo claimed by each country (Indonesia: 72.5% of land area, Malaysia: 26.7% and Brunei: 0.8%) and summed to give GDP per capita, PPP and SPI estimate for Borneo.

That is, using Equation 1, the SPI for Borneo was calculated as:

$SPI^{BORNEO} = 93.14(195376/735738) + 77.57(534683/735738) + 99.89(5677/735738) = 81.88$

Population density was calculated for each country by dividing total population by land area, in $km^2$, assuming a Behrmann Equal area projection. Where a country was shared by multiple sovereign nations, population was allocated to each unique combination of WGSRPD unit and sovereign nation proportional to the amount of land area claimed. The population of each country was then calculated as the sum of population in each of the combinations following Equation 2:

Eq. 2: $$p^{BOT} = \sum_{i=1}^{n} p_i^{SOV} \left(\frac{a_i^{UNIT}}{a^{SOV}}\right)$$

Where $\boldsymbol{p}^{BOT}$ was the estimated population of the country, $\boldsymbol{p}^{SOV}$ was the population of the sovereign nation, $\boldsymbol{a}^{UNIT}$ was the area of the sovereign nation within the country, and $\boldsymbol{a}^{SOV}$ was the total area of the sovereign nation.

That is, using Equation 2, the population for Borneo was calculated as:

$$p^{BOR} = 3.15 \times 10^7 \left(\frac{195376}{327555}\right) + 2.68 \times 10^8 \left(\frac{534683}{1.87 \times 10^6}\right) + 428962 \left(\frac{5677}{5677}\right)$$
$$= 9.56 \times 10^7$$

This population is then divided by the land area of Borneo to give the population density in people per $km^2$:

$$pd^{BOR} = \frac{9.56 \times 10^7}{735738} = 130$$

Although population density is a single value for each country, we recognise that density will vary across space and human influence on endemic flora may be larger in some regions relative to others, such as capital cities and coastlines.

### 6. Climate change and deforestation threat in countries

Exposure to climate change was assessed using gridded data on current (1979-2013) and future (2070) mean annual temperature (MAT; °C) under representative concentration pathway 8.5 which was accessed from the CHELSA portal[13] at a 30 arc-second resolution. A single, future MAT projection for 2070 was created by calculating the median across seven global climate models (CESM1-BGC, MPI-ESM-MR, MIROC5, CMCC-CM, CESM1-CAM5, IPSL-CM5A-MR, FIO-ESM). Taking the median of a suite of GCM projections represents variation in spatial patterns of future temperature[14]. Anomalies were calculated between current and future MAT projections (Fig. S11A) and the average anomaly across each country was calculated to quantify potential exposure to climate change (Fig. S11B).

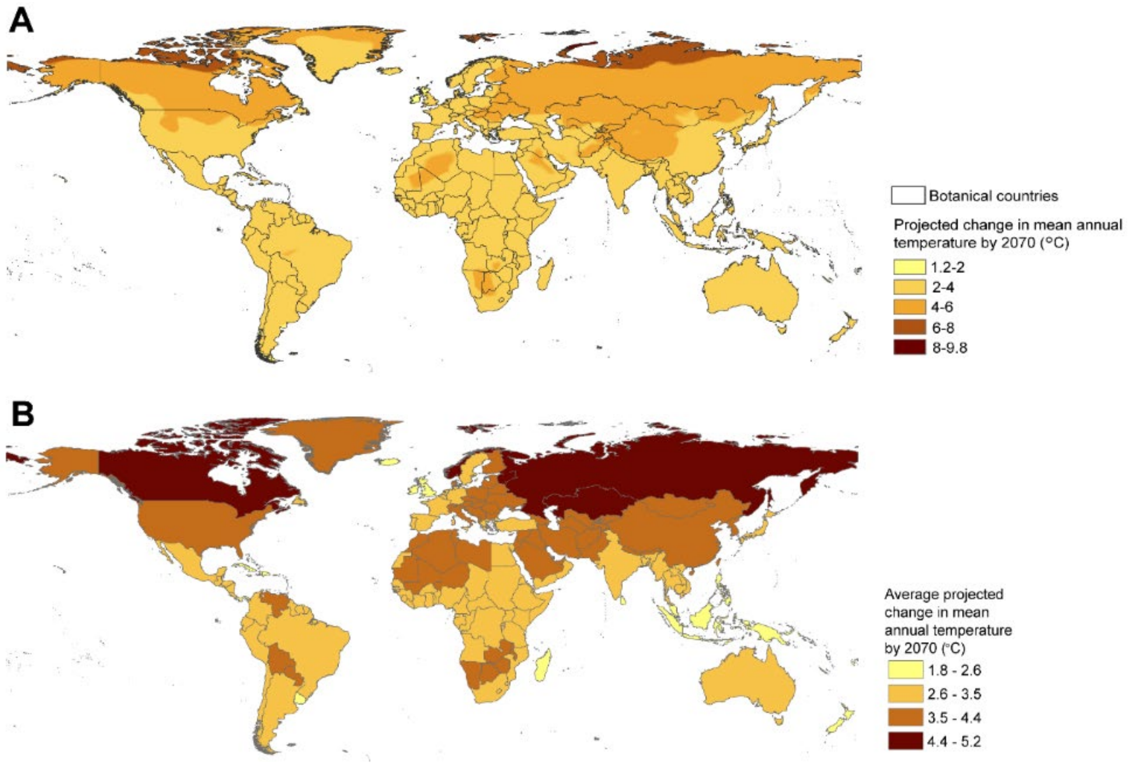


**Figure S11.** (A) Exposure to changes in mean annual temperature (MAT) across the countries of the world produced by calculating anomalies between current (1979-2013) MAT and future projections of MAT from seven Global Climate Models. (B) Average values of MAT anomalies in each country used in analyses.

Gridded data on permanent deforestation was accessed from the Supplementary Materials of [15,16]. Permanent deforestation was defined as land use conversion that prevents subsequent forest regrowth[15] and was calculated for grid cells with a canopy cover >30%. We recognise that a threshold of >30% canopy cover may exclude some areas of the globe known to be experiencing significant deforestation, such as Madagascar[17].

Two drivers of forest loss were combined spatially to identify areas of permanent deforestation: commodity-driven deforestation (category 1 in[15]) and urbanization (category 5 in[15]) (Fig. S12). This composite map is based on classification of remotely sensed imagery used to identify all 10 km x 10 km grid cells (Berhmann equal-area projection) where drivers of permanent deforestation were the most likely cause of forest disturbance since 2000. The percentage of deforested area in each country was used as an indicator of threat.

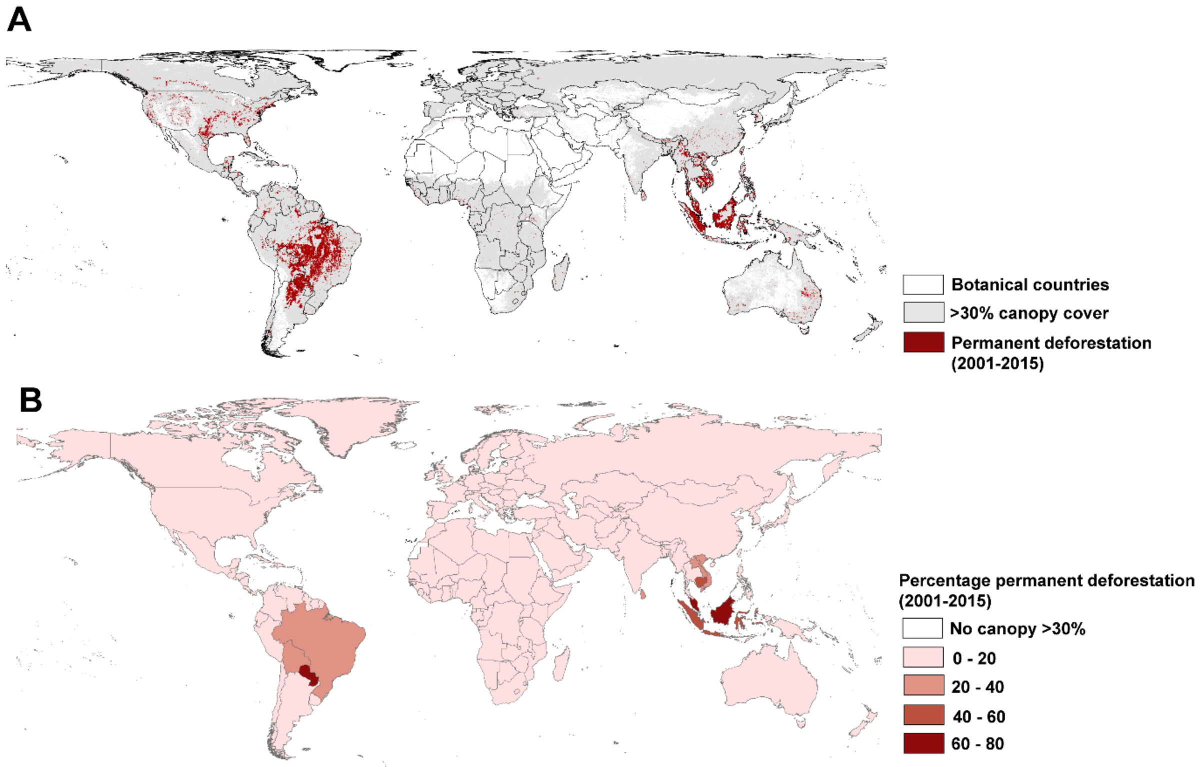


**Figure S12**. (A) Areas with canopy cover >30% experiencing permanent deforestation between 2001 and 2015 (B) Percentage of a country with >30% canopy cover subject to permanent deforestation between 2011 and 2015.

### 7. Statistical analyses

Data and code used to complete statistical analyses are available in Dataset S6.

The association between the proportion of endemic plant species with a threat assessment and the total number of endemic plant species in countries was quantified using Kendall's rank correlation coefficient (Kendall's $\tau$) using the *DescTools*[18] in R[19]. A 95% confidence interval for $\tau$ was calculated via bootstrapping using the function *KendallTauA*. Tests were considered significant at an alpha level of 0.05.

Generalised additive mixed models (GAMMs) were used to quantify the relationship between the proportion of endemic species assessed in countries; a set of economic, demographic and threat predictor variables (*i.e.*, GDP per capita, PPP, population density, SPI, climate change exposure and permanent deforestation; Table S1); and the spatial distance between countries (latitude/longitude of country centroids). GAMMs were chosen as they allow the effect of non-linear spatial trends to be combined with linear predictors. The latitude and longitude coordinates at the centroid of each country were added to GAMMs as a two-dimensional spherical spline smoother, *f*(lat,lon). GAMMs were fit in R using package *gamm4*[20] and visualised using the package *gratia*[21].

**Table S1.** Descriptive statistics of linear predictor variables used in generalized additive mixed model used to predict relationships with proportion of endemic plant species with an extinction risk assessment in countries globally (*n* = 177).

| **Predictor** | ***N*** | **Mean** | **SD** | **Minimum** | **Maximum** |
|---|---|---|---|---|---|
| GDP per capita ($USD) | 177 | 13106.02 | 17831.36 | 275.43 | 82838.93 |
| Purchasing Power Parity ($USD) | 177 | 397.08 | 1369.78 | 0.14 | 11427.68 |
| Population density (individuals/ $km^2$) | 177 | 139.45 | 299.98 | 2.03 | 3655.42 |
| Species Protection Index (0-100) | 153 | 71.55 | 28.54 | 2.61 | 100.00 |
| Climate change exposure (°C) | 169 | 3.19 | 0.65 | 1.75 | 5.22 |
| Permanent deforestation 2000-2015 (%) | 165 | 4.68 | 11.57 | 0.00 | 74.17 |

$USD = United States dollars

All variables, except climate change exposure, were transformed prior to analysis. GDP per capita, PPP, and population density were log-transformed to correct skew and equalize variance relative to the mean. The proportional variables SPI and endemics assessed (the response) were constrained to values between 0-100 and had reasonably small numbers of observations on the zero or hundred bounds (*n* = 10 of 177 observations of SPI were 100; *n* = 13 of 177 observations of endemics assessed were 0). Therefore, these variables were logit transformed to reduce overdispersion following Equation 3:

Eq. 3: $$Yi = \log((Pi + x)/(1-Pi + x))$$

where, for the response variable, *x* was the value closest to the zero bound/2 (i.e., 0.0035) and, for SPI, *x* was the value closest to the hundred bound/2 (i.e., 0.005)[22]. Permanent deforestation was also bounded (0-100), though had a large number of true xero values (*n* = 64 of 165 observations) and logit transformation did not adequately address overdispersion. Therefore, permanent deforestation was analysed as a binary factor (TRUE/FALSE) in all analyses.

We fit three alternative GAMMs to all predictors and partitioned variance in the residuals using the *ecospat*[23] package:

(1) a ***full*** model including a spatial smoother, $f(\text{lat,lon})$, along with all linear predictors, $x_i$, and their associated coefficients, $\beta_i$,
$$y_i = \beta_0 + x_1\beta_1 + x_2\beta_2 + x_3\beta_3 + x_4\beta_4 + x_5\beta_5 + x_6\beta_6 + f(\text{lat,lon}) + \varepsilon_i,\ \varepsilon_i \sim N(0,\sigma^2)$$

(2) a ***spatial*** model including only the smoother
$$y_i = \beta_0 + f(\text{lat,lon}) + \varepsilon_i,\ \varepsilon_i \sim N(0,\sigma^2)$$

(3) an ***aspatial*** model including only the linear predictors
$$y_i = \beta_0 + x_1\beta_1 + x_2\beta_2 + x_3\beta_3 + x_4\beta_4 + x_5\beta_5 + x_6\beta_6 + \varepsilon_i,\ \varepsilon_i \sim N(0,\sigma^2)$$

We assumed a Gaussian distribution and identity link for all GAMMs. Several routine diagnostic plots confirmed that GAMM assumptions were reasonably met for all models, including checks for normality, equal variance of residuals and linearity (Fig. S13-15).

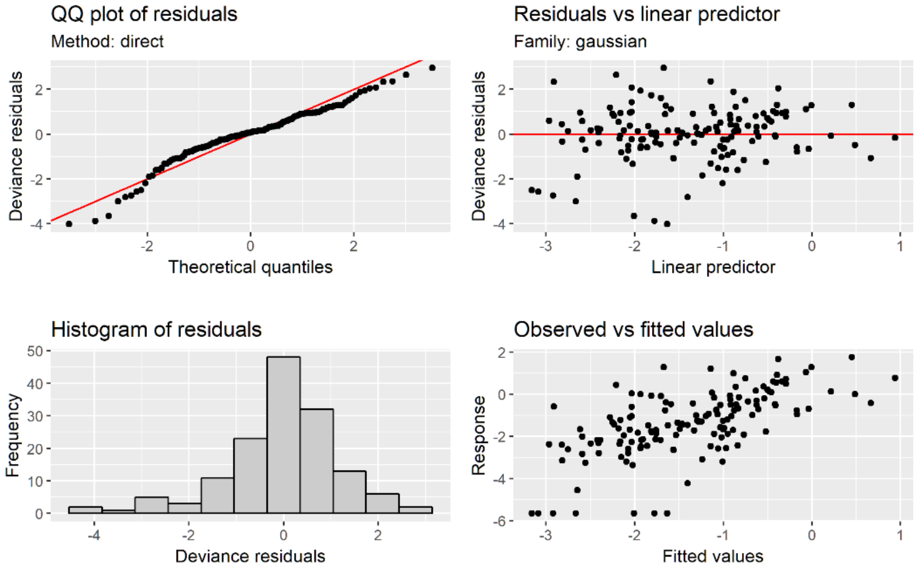


**Figure S13.** Diagnostic plots for the *full* generalised additive mixed model (GAMM) including six linearised predictors (GDP per capita, PPP, population density, SPI, climate change exposure and permanent deforestation) and a two-dimensional spherical spline smoother to account for non-linear spatial trends. $y = \beta_0 + x_1\beta_1 + x_2\beta_2 + x_3\beta_3 + x_4\beta_4 + x_5\beta_5 + x_6\beta_6 + f(x_7) + \varepsilon, \varepsilon \sim N(0,\sigma 2)$

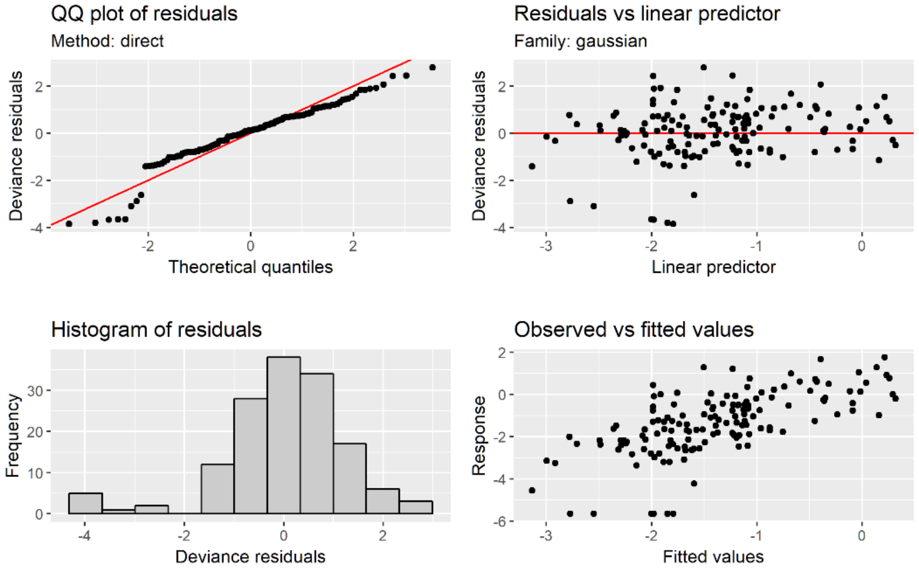


**Figure S14.** Diagnostic plots for the *spatial* generalised additive mixed model (GAMM) based on a two-dimensional spherical spline smoother. $yi = \beta_0 + f(\text{lat,lon}) + \varepsilon i, \varepsilon i \sim N(0,\sigma^2)$

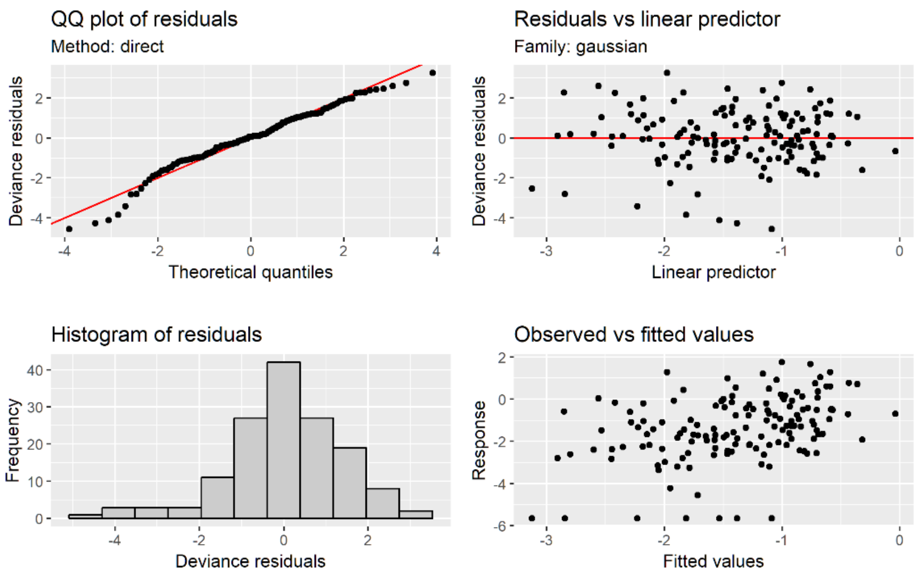


**Figure S15.** Diagnostic plots for the *aspatial* generalised additive mixed model (GAMM) including six linearised predictors (GDP per capita, PPP, population density, SPI, climate change exposure and permanent deforestation). $y = \beta_0 + x_1\beta_1 + x_2\beta_2 + x_3\beta_3 + x_4\beta_4 + x_5\beta_5 + x_6\beta_6 + \varepsilon$, $\varepsilon \sim N(0,\sigma 2)$

Diagnostic residual plots indicated that standard methods for accounting for overdispersion were not appropriate when proportion of endemics was analysed as a binomial variable[22].

Residual variance was partitioned into components related to the unique contributions of our predictors and space, as well as their covariance, following methods in[24]. Hypothesis tests on the Moran's I correlation statistic for each model (i.e., *z*-scores and their associated *p*-values) were used to quantify the independence of linear predictors and space in all models. Moran's I is a unitless value that ranges between -1 (strong negative correlation between the response and distance, here degrees of latitude) to 1 (strong positive correlation)[25]. Finally, we mapped patterns of residual variance in the smoothing spline to visually inspect the spatial structure of the full model.

### 8. Results

A significant association was identified between the proportion of endemic species in a country with an extinction risk assessment and the total number of endemic plants per country (Kendall's $\tau = 0.17$ (CI: 0.07-0.27); $p = 0.001$). Although significant, this association was not strong ($\tau = 0.17$) meaning that the magnitude of plant endemism is of little value in predicting a country's progress toward Target 2 of the Global Strategy for Plant Conservation.

Our *full* GAMM model indicated that the proportion of endemic plant species with an extinction risk assessment showed no significant relationships ($p > 0.05$) with any of the six hypothesised predictors (GDP per capita, PPP, population density, SPI, climate change exposure, presence/absence of permanent deforestation) after accounting for highly significant broad-scale spatial trends ($p < 0.001$) captured by the spline smoother (Table S2).

**Table S2.** Parameter estimates and approximate significance level of linear predictors and a spline smoother on proportion of endemic plant species with an extinction risk assessment in countries using a generalized additive mixed model (GAMM). The spherical spline smoother is calculated using great-circle distances between the latitude-longitude centroids of the countries.

| **Parameters** | **Estimates** | **St. error** | ***t*-statistic** | ***P*-value** |
|---|---|---|---|---|
| Intercept | -4.17 | 1.92 | -2.18 | 0.03* |
| GDP per capita | 0.26 | 0.14 | 1.91 | 0.06 |
| Purchasing Power Parity | -0.07 | 0.05 | -1.35 | 0.18 |
| Population density | 0.21 | 0.12 | 1.78 | 0.08 |
| Species Protection Index | 0.04 | 0.05 | 0.78 | 0.44 |
| Climate change exposure | -0.18 | 0.31 | -0.58 | 0.57 |
| Permanent deforestation TRUE | 0.62 | 0.33 | 1.87 | 0.06 |
| | **EDF[1]** | **Ref. DF[2]** | ***F*-statistic** | ***P*-value** |
| **Smoothing function** | | | | |
| Latitude/longitude | 15.76 | 49 | 0.766 | <0.001* |

EDF = effective degree of freedom; 2. Ref. DF = reference number of degrees of freedom

Moran's I statistics (calculated using inverse great-circle distances between centroids) indicated that the inclusion of the spherical spline smoother in the *full* GAMM adequately reduced spatial autocorrelation in the residuals, relative to the aspatial model (Table S3).

**Table S3.** Hypothesis tests on *z*-scores of Moran's I (observed and expected values), standard deviation and P-values across three GAMMs: *full* (all linearised predictors and a spline smoother); *Spatial* (spline smoother only); *Aspatial* (linearised predictors only).

| **GAMM** | **Moran's I (observed)** | **Moran's I (expected)** | **SD** | ***P*-value** |
|---|---|---|---|---|
| Full | -0.02 | -0.01 | 0.01 | 0.20 |
| Spatial | -0.03 | -0.01 | 0.01 | 0.04 |
| Aspatial | 0.04 | -0.01 | 0.01 | <0.0001 |

Our *full* model including all predictor variables and the spatial smoother accounted for 67.5% of the variance, but variance partitioning indicated that none of variance could be confidently attributed to our economic, demographic or threat predictors after accounting for broad-scale spatial trends (-0.02% variance). Partitioning showed that 32.3% of variance in the *full* model reflected spatial variation unrelated to our target variables, and 35.4% reflects covariance between predictor variables and space. Partial residual plots show little predictive structure to the relationship between endemic assessment and each variable (Fig. S16) and mapping of the spatial smoother indicate strong structure in the residuals (Fig. S17). Therefore – despite our hypotheses about the importance of economic wealth, threat exposure, human demography or species protection on efforts to assess extinction risk in plants – our suite of variables does not predict the progress of countries toward Target 2 of the Global Strategy for Plant Conservation.

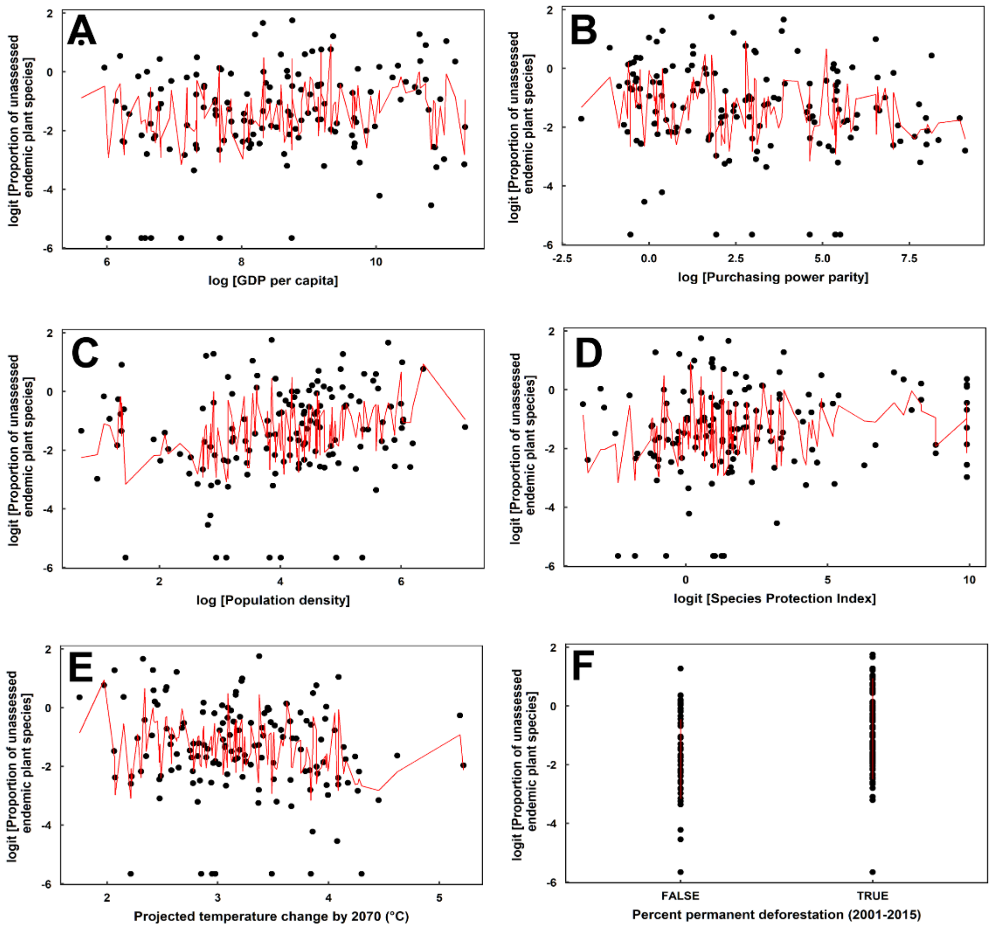


**Figure S16.** Partial residual plots for the *full* generalised additive mixed model (GAMM) including six linearised predictors (GDP per capita, PPP, population density, SPI, climate change exposure and permanent deforestation) and a two-dimensional spherical spline smoother to account for non-linear spatial trends.

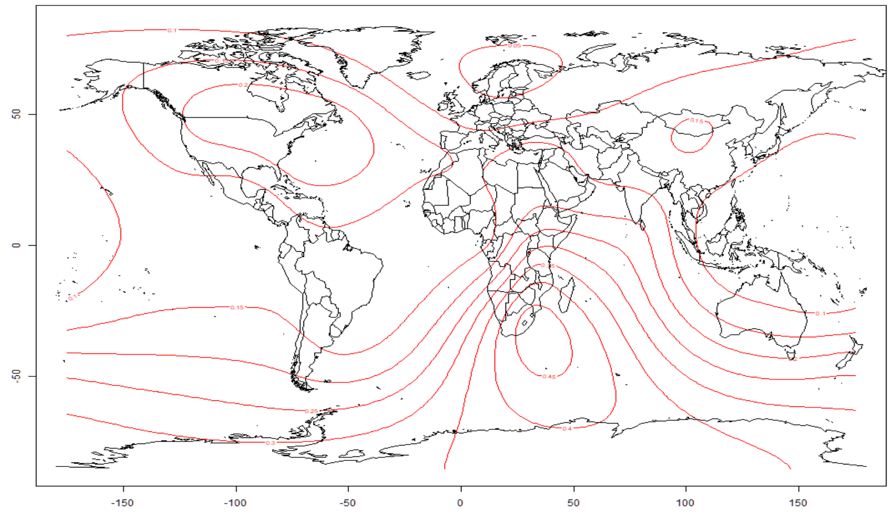


**Figure S17.** Map of the spatial spline smoother *f*(lat,long) across countries. Closer contour lines indicate areas of stronger spatial structure in the residuals of the *full* GAMM model.
